# GDF15 contributes to inflammasome-associated excessive mechanoresponses of hyperlipidemic PdL fibroblasts

**DOI:** 10.64898/2026.08.30.748125

**Authors:** Martin Baumbach, Annamaria Manzolillo, Faezeh Ghazvini Zadegan, Raushan Yeskendirova, Annika Döding, Christoph-Ludwig Hennig, Ulrike Schulze-Späte, Judit Symmank, Collin Jacobs

**Affiliations:** Department of Orthodontics, Jena University Hospital, Jena Germany; Section of Geriodontics, Department of Conservative Dentistry and Periodontics, Jena University Hospital, Jena, Germany; Department of Orthodontics, University Hospital Düsseldorf, Düsseldorf, Germany

**Keywords:** periodontal ligament fibroblasts, orthodontic tooth movement, palmitic acid, pyroptosis, GDF15

## Abstract

Orthodontic tooth movement relies on a tightly regulated pro-inflammatory and pro-resorptive mechanoresponse of local periodontal ligament fibroblasts (PdLFs). Dysregulation is linked to complications such as root resorption and tooth loss. Hyperlipidemic conditions promote excessive PdL mechanoresponses, with growth differentiation factor 15 (GDF15) acting as potential regulator. This study examined the contribution of the inflammasome/pyroptosis pathway as underlying mechanism for dysregulated mechanoresponses. Human PdLFs were treated with palmitic acid (PA) or oleic acid (OA) for six days before 24 hours of compressive loading. PA increased CASP1, CASP4, and CASP3 activity, secretion of IL-1β, IL-18, and HMGB1, and LDH release. Pharmacological blockade and siRNA-mediated knockdown of inflammasome– and pyroptosis-related targets revealed that NLRP3, CASP1, CASP4, and GSDMD partially contributed to monocyte and osteoclast overactivation. Silencing PA-increased GDF15, partially normalized the phenotype, at least in part by inflammasome/pyroptosis regulation. GDF15 acted through extracellular, and a nuclear signaling route, each accounting partially to this phenotype. Together, GDF15 partially regulates the PA-induced, pyroptosis-associated overactivated mechanoresponse alongside pyroptosis-independent mechanisms suggesting it as an interesting target for potential clinical interventions.

## Introduction

Orthodontic tooth movement depends on a coordinated tissue and bone remodeling driven by mechanical forces acting specifically on connective tissue surrounding the teeth, the periodontal ligament (PdL) [1]. Fibroblasts are the most abundant cell type in the PdL converting mechanical stimuli into inflammatory responses by secreting of pro-inflammatory cytokines including interleukin 6 (IL-6), IL-1β, tumor necrosis factor-α (TNF-α), and cyclooxygenase 2 (COX2). PdL fibroblasts further secrete receptor activator of nuclear factor kappa-B ligand (RANKL) and osteoprotegerin (OPG), which are relevant regulators in bone remodeling [2]. Receptor activator of nuclear factor kappa-B (RANK) is expressed by osteoclast precursors and binding of its ligand RANKL promote their differentiation. OPG acts as a decoy receptor, binding RANKL and preventing this interaction. Compressive force increases this ratio promoting osteoclast differentiation and resorption on the compression side, which enables tooth movement [3]. An excessive or dysregulated activation of this mechanoresponse potentially increases the risk of orthodontic treatment-associated complications, including root resorption and tooth loss [4, 5]. Patient-related and –specific factors that shift this mechanical response toward a hyperinflammatory state are therefore of particular clinical interest.

Obesity-associated hyperlipidemia induces hyperinflammatory responses in mechanically-stressed connective tissue, with the underlying triggers and mechanisms appearing highly multifactorial in regard to fatty acid and tissue specificity [6, 7]. In line with this, hyperlipidemic conditioning with the saturated palmitic acid (PA), but not with the unsaturated oleic acid (OA), has previously been demonstrated to overactivate the inflammatory mechanoresponse of PdL fibroblasts [8–10]. The activation of cell death pathways is considered a possible underlying mechanism of hyperinflammatory tissue responses. In this regard, saturated, but not unsaturated, fatty acids were reported to prime and activate the NLRP3 inflammasome leading to pyroptosis [11, 12], also specifically in PdL cells [13]. Pyroptosis is a programmed necrotic cell death that ultimately results from the activation of the inflammasome signaling pathway, whether via the canonical or non-canonical route [14]. In the canonical pathway, NLR family pyrin domain containing 3 (NLRP3) assembles with the adaptor protein ASC to activate caspase-1 (CASP1), which cleaves gasdermin D (GSDMD) and the pro-inflammatory cytokines IL-1β and IL-18 into their active forms. Pore-formation by GSDMD allowing the release of these cytokines along with high mobility group box 1 (HMGB1) and lactate dehydrogenase (LDH), triggering a potent inflammatory response. A non-canonical route exists through CASP4 and CASP5, which can be activated independently of NLRP3. Force-induced inflammasome activation has been shown to regulate orthodontic tooth movement, with CASP1-dependent PdL cell pyroptosis contributing to osteoclastogenesis and alveolar bone remodeling [15].

It has been consistently demonstrated that growth differentiation factor 15 (GDF15) modulates the mechanical response of PdL fibroblasts in a pro-inflammatory and pro-resorptive manner [16–21]. As part of the transforming growth factor beta (TGF-β)/bone morphogenic protein (BMP) superfamily, GDF15 is weakly expressed under physiological conditions and induced by cellular stress with upregulated levels under hyperinflammatory conditions in the PdL [17, 19, 21].

GDF15 is synthesized as a precursor and can be cleaved into a mature, secreted homodimer or retained as an uncleaved form with distinct intracellular, nuclear, and extracellular matrix-bound localizations, each associated with a different mode of action [22, 23]. GFRAL, together with the coreceptor RET, is the best-characterized GDF15 receptor, though its expression is restricted to a subset of neurons in the brainstem [24]. In PdL fibroblasts, expression of peripherally relevant GDF15 receptors of the activin receptor like kinase (ALK) family, specifically ALK1, ALK2, and ALK5, was recently demonstrated [18].

Circulating GDF15 levels are elevated in patients with hyperlipidemia and correlate with atherogenic lipid measures [25]. However, a direct link between GDF15 and the inflammasome/pyroptosis pathway has not been established. This study aimed to investigated whether palmitic acid activates this pathway in PdL fibroblasts under compressive force, and whether GDF15 contributes through its extracellular and intracrine signaling routes. Identifying this connection might help clarify targets for limiting hyperinflammatory, treatment-associated risks in orthodontic patients with hyperlipidemia.

## Material and Methods

### Cell culture

Human periodontal ligament fibroblasts (hPdLFs; CC-7049, Lonza, Basel, Switzerland) were expanded in high-glucose Dulbecco’s modified Eagle medium (DMEM with 4.5 g/L glucose; Thermo Fisher Scientific, Carlsbad, CA, USA). The medium was supplemented with 10% heat-inactivated fetal bovine serum (FBS), 100 U/mL penicillin, 100 µg/mL streptomycin, and 50 mg/L L-ascorbic acid. Cultures were maintained at 37°C under humidified 5% CO₂ conditions. At approximately 75% confluence, cells were detached using 0.05% trypsin/EDTA and reseeded as required. Experiments were performed with hPdLFs between passages four to eight.

THP1 monocytic cells (DSMZ, Braunschweig, Germany) were propagated in RPMI 1640 supplemented with 10% FBS, 100 U/mL penicillin, and 100 µg/mL streptomycin. Cultures were maintained at 37°C and 5% CO₂ and routinely passaged to prevent excessive cell density.

### Fatty acid exposure

For fatty acid conditioning, hPdLFs were exposed to either palmitic acid (PA) or oleic acid (OA) at a final concentration of 200 µM based on previously established conditions [8, 10]. Briefly, fatty acids were first solubilized at 70°C in sterile water containing 50 mM NaOH and subsequently complexed with pre-warmed BSA. The resulting fatty acid-BSA complexes were diluted into standard fibroblast culture medium. Control cultures received medium containing the corresponding BSA concentration of 0.66% without fatty acid. Cells were maintained under these conditions for six days before subsequent experimental stimulation.

### siRNA-mediated knockdown

For transient gene silencing, hPdLFs were transfected at approximately 75% confluence as previously described [21, 26]. Briefly, target-specific siRNA (all Santa Cruz Biotechnology, Dallas, Texas, USA) was used at a final concentration of 50 nM and transfected using Lipofectamine™ 2000 (Thermo Fisher Scientific) in Opti-MEM™ I Reduced Serum Medium (Thermo Fisher Scientific) supplemented with 100 U/mL penicillin and 100 µg/mL streptomycin. After 5 h of transfection, medium was replaced with regular PdL fibroblast culture medium. A non-targeting siRNA was included as the corresponding negative control. Cells were subsequently subjected to the experimental treatments specified for the individual assays.

### Application of inhibitors

Pharmacological inhibitors were used to interfere with inflammasome/pyroptotic signaling, activin receptor-like kinase (ALK) signaling, and nuclear protein export. The NLRP3 inflammasome was inhibited using 5 µM MCC950, whereas 25 µM VX-765 was applied to inhibit CASP1/4/5 activity. To discriminate between different branches of ALK signaling, cells were treated with 0.5 µM MU1700, targeting predominantly ALK1/ALK2, or 1 µM TP-008, an ALK4/ALK5 inhibitor. All inhibitors were applied in PA-containing media 24 hours before compressive force application. Nuclear export was inhibited with 10 nM leptomycin B (LMB), applied for 90 min prior compressive force application. Since LMB significantly affected cell viability in preliminary experiments (data not shown), culture medium was changed to PA-containing media without LMB before compression was started. Corresponding vehicle-treated cells were included as controls.

### Compressive force application

Mechanical compression was generated by positioning sterile polytetrafluoroethylene plates directly onto confluent hPdLF monolayers comparable to previous protocols using glass plates [21, 26]. The plates produced a static pressure of 2 g/cm², which was maintained for 24 hours at 37°C and 5% CO₂. Parallel cultures maintained under identical incubation conditions without mechanical loading served as controls.

### Quantitative expression analysis

The underlying RNA and qPCR workflow follows previously established protocols [8, 26]. Briefly, total RNA was extracted from hPdLF cultures using TRIzol™ Reagent (Thermo Fisher Scientific). Following extraction, RNA was further purified with the RNA Clean & Concentrator-5 Kit (Zymo Research, Freiburg, Germany). RNA concentration and purity were assessed spectrophotometrically before reverse transcription. Complementary DNA was generated using SuperScript™ IV Reverse Transcriptase together with oligo(dT)18 primers. Quantitative real-time PCR was carried out with Luminaris Color HiGreen qPCR Master Mix (Thermo Fisher Scientific) using a qTOWER³ real-time PCR system (Analytik Jena, Jena, Germany). Primer pairs were evaluated for amplification specificity and efficiency prior to experimental use. *RPL22* and *TBP* served as internal reference genes. Relative transcript abundance was determined using an efficiency-corrected ΔΔCT approach. Primer sequences and information are provided in Table 1.

**Table 1:**
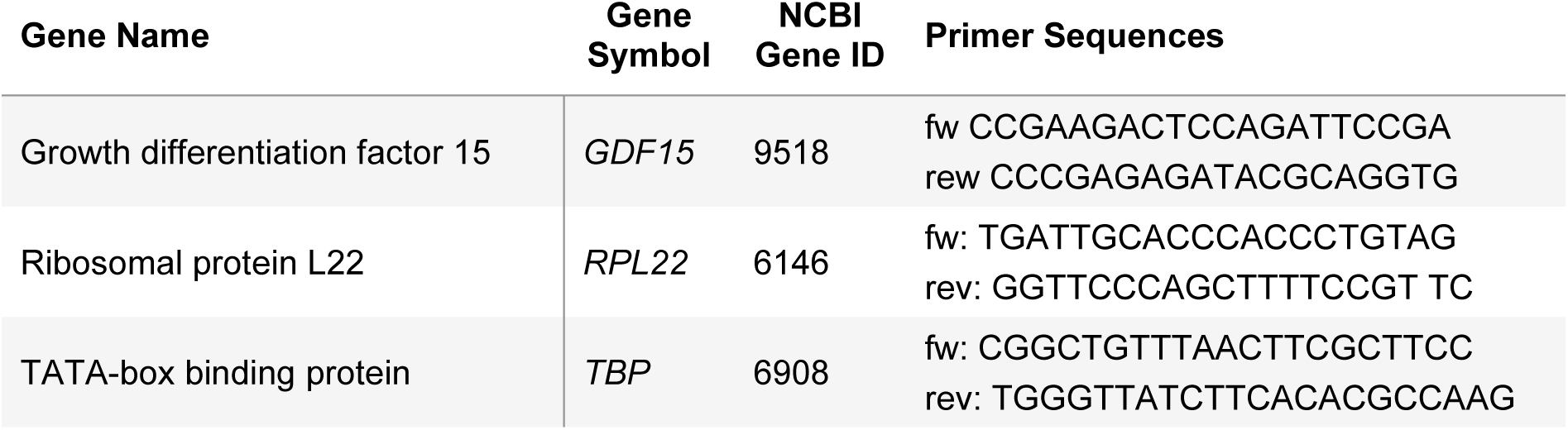
Forward (fw) and reverse (rev) primer for quantitative expression analysis. Gene-specific exon-spanning primers are indicated in 5’-3’ direction.

### Immunofluorescent staining

The procedure for immunofluorescent stainings was adapted from previously established protocols [18, 26]. For immunofluorescence analysis, cells were grown on glass coverslips and fixed with 4% paraformaldehyde for 10 min. Following fixation, membranes were permeabilized using 0.1% Triton X-100 in phosphate-buffered saline (PBS). Samples were subsequently incubated in blocking solution (PBS/0.1% Triton X-100 with 4% BSA) to reduce nonspecific antibody binding. The primary antibody targeting GDF15 (Thermo Fisher Scientific; MA531346) was diluted 1:200 in blocking solution and incubated with the samples for 3 h at room temperature. After washing, a Cy5-conjugated anti mouse secondary antibody was applied for 45 min. Nuclear DNA was visualized with DAPI diluted 1:10,000 in PBS. Coverslips were finally mounted using Mowiol® 4-88.

### Enzyme-Linked Immunosorbent Assay

Secreted protein levels were determined in cell culture supernatants using the commercially available enzyme-linked immunosorbent assay kits for GDF15 (Thermo Fisher Scientific; BMS2258), IL-1β (Abcam, Cambridge, UK ab214025), IL-6 (Abcam, ab178013), IL-18 (Abcam; ab215539) and HMGB1 (Thermo Fisher Scientific; EEL047). Supernatants were harvested after completion of the respective stimulation period, cleared of cellular material by centrifugation at 1000 rpm for 5 min, and stored at −20°C until analysis. Samples were measured in technical duplicates in accordance with the instructions supplied by the respective manufacturer. Concentrations were calculated from assay-specific standard curves.

### LDH Assay

Loss of cell membrane integrity was evaluated by measuring lactate dehydrogenase (LDH) released into the extracellular medium. Following experimental stimulation, culture supernatants were collected and analyzed with the Lactate Dehydrogenase (LDH) Colorimetric Activity Kit (Thermo Fisher Scientific; EEA013) according to the manufacturer’s protocol. Each sample was measured in technical duplicate.

### Caspase activity assay

Caspase activity was assessed using the Caspase Family Colorimetric Substrate Kit II Plus (Abcam; ab102488) according to the manufacturer’s instructions. Activities of CASP1, CASP3, CASP4, CASP5, CASP8, and CASP9 were quantified using the corresponding p-nitroaniline (pNA)-conjugated substrates. Absorbance was measured at 405 nm, and background-corrected values were normalized to the corresponding control condition.

### THP1 adhesion assay

The inflammatory potential of stimulated hPdLF was assessed using the adhesion of THP1 monocytes as previously established as functional readout [21, 26]. THP1 cells were first labelled with CellTracker™ Green CMFDA (Thermo Fisher Scientific). For each condition, 2.5 × 10⁴ labelled THP1 cells were cocultured with stimulated hPdLFs grown on glass coverslips in 24-Well plates. After 30 min at 37°C, cultures were washed with pre-warmed PBS to remove non-adherent THP1 cells. Cells remaining attached to the culture surface were fixed with pre-warmed 4% paraformaldehyde, stained with DAPI and subsequently analyzed by microscopy.

### Osteoclast activation assay

THP1 cells were used as an osteoclast precursor model to assess the osteoclastogenic activity of hPdLF-conditioned media as previously established [18, 19, 26] Briefly, monocytic THP1 cells were initially exposed to 100 ng/mL phorbol 12-myristate 13-acetate (PMA; Merck Millipore) for 48 h to induce differentiation toward an adherent macrophage-like phenotype. After this initial differentiation step, cells were maintained for further six days in a 1:1 mixture of fresh THP1 culture medium and conditioned medium collected from the respective hPdLF treatment groups. Finally, cells were fixed using pre-warmed 4% paraformaldehyde and processed for tartrate-resistant acid phosphatase staining.

### Tartrate-resistant acid phosphatase (TRAP) staining

Osteoclast-like differentiation was visualized by histochemical detection of tartrate-resistant acid phosphatase activity as described previously [18, 26]. Briefly, fixed THP1 cultures were incubated for 60 min at 37°C in a staining solution containing 0.1 mg/mL Naphthol AS-MX phosphate, 0.5 mg/mL Fast Red Violet LB salt, and 1% N,N-dimethylformamide. The substrate solution was prepared in 50 mM sodium acetate containing 50 mM tartrate and 0.1% acetic acid. Following TRAP staining, nucleic acids were counterstained with SYTO for 5 min. Samples were washed with PBS and imaged immediately. Multinucleated cells displaying positive TRAP staining were classified as osteoclast-like cells and quantified relative to the corresponding experimental control.

### Microscopy and Image Analysis

Imaging of immunofluorescent stainings and fluorescently labelled THP1 cells was performed using a LSM 880 (Carl Zeiss AG, Jena, Germany TRAP/SYTO-stained cultures were documented using a Primovert microscope (Carl Zeiss AG). For comparisons between treatment groups, microscope and camera settings were kept unchanged within each experimental series. Image quantification was carried out using Fiji/ImageJ. Fluorescence intensity was quantified as mean gray value after correction for local background. Where applicable, measurements were normalized to the corresponding untreated or vehicle-treated control. THP1 activation was quantified from the number of adherent CMFDA-positive cells to all DAPI-positive cells, which include hPdLFs. Osteoclast differentiation was assessed by counting multinucleated TRAP-positive cells in TRAP/SYTO-stained cultures.

### Statistics

Statistical analyses were performed using GraphPad Prism 8 (GraphPad Software, Boston, MA, USA). For experimental designs containing more than two groups or factors, one-way or two-way analysis of variance (ANOVA) was applied, followed by Tukey’s multiple-comparisons test. Unless specified otherwise, results are shown as mean ± standard error of the mean (SEM) from at least three biologically independent experiments. A p-value less than 0.05 was considered statistically significant.

## Results

### Palmitic acid activates the inflammasome/pyroptosis pathway

Palmitic acid (PA) has been reported to induce GSDMD-mediated pyroptosis in periodontal ligament cells [13]. To determine whether this signaling pathway could be contributing to hyperinflammatory mechanoresponses observed in obesity-associated hyperlipidemic conditions [9], we analyzed human PdL fibroblasts exposed to PA for six days prior to application of compressive force (CF). Oleic acid was additionally included to identify fatty acid-specific responses rather than those caused by excessive lipid stimulation. BSA was used as a control condition. Caspase activity assays revealed a significant activation of inflammasome-associated caspases CASP1 and CASP4 as well as apoptosis-related CASP3 in PA+CF cultures compared to compressed BSA controls (**Fig. 1a**). Secretion of pyroptosis-associated cytokines including IL-1β and IL-18, and of the membrane-rupture marker HMGB1, was significantly elevated in PA+CF relative to BSA+CF (**Fig. 1b**). This was in line with an increased LDH release in PA+CF (**Fig. 1c**). OA+CF demonstrated similar results to BSA+CF for all three measured parameters. Together, these data confirm PA-specific inflammasome/pyroptosis pathway activation in hPdLFs under compressive force.

**Figure 1:**
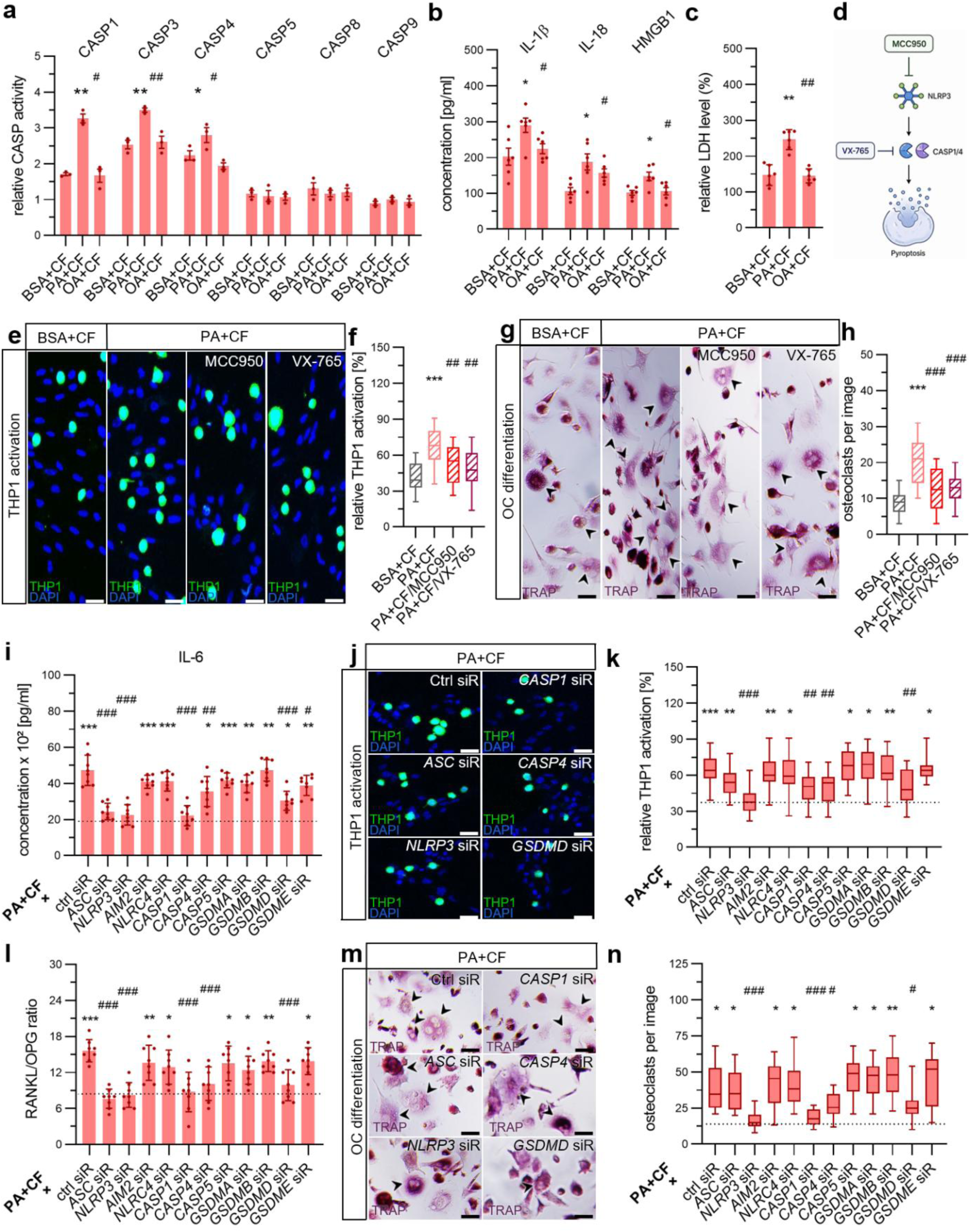
Palmitic acid activates the inflammasome/pyroptosis pathway in compressed PdL fibroblasts, which is functionally required for the hyperinflammatory mechanoresponse. Prior compressive force (CF) application of 24 hours, human periodontal ligament fibroblasts (hPdLFs) were treated with palmitic acid (PA) or oleic acid (OA) with bovine serum albumin (BSA)-treated cells as control (a) Activity of CASP1, CASP3, CASP4, CASP5, CASP8, and CASP9 in compressed PA– and OA-treated hPdLFs relative to an unstressed BSA control. (b) Secretion of IL-1β, IL-18, and HMGB1 by compressed PA– and OA-treated hPdLFs. (c) LDH release in medium supernatant of compressed PA– and OA-treated hPdLFs. (d) Schematic of pharmacological pyroptosis inhibition using MCC950 targeting NLRP3 or VX-765 targeting CASP1/4/5. (e, f) Activation of fluorescently labeled monocytic THP1 cells (green, e) after coculturing with stressed hPdlFs analyzed as percentage of THP1 cells to all cells per condition. Cell nuclei were labeled by DAPI. (g, h) Osteoclast (OC, magenta, g) activation in compressed PA and OA-treated hPdLFs following VX-765 or MCC950 treatment analyzed as multinucleated OC per image. (i) IL-6 secretion following siRNA-mediated knockdown of inflammasome– and pyroptosis-associated targets in compressed PA and OA-treated hPdLFs. (j, k) THP1 activation in siRNA-treated hPdLFs. (l) RANKL/OPG ratio following siRNA knockdown. (m, n) Osteoclast activation following siRNA knockdown. One-way ANOVA with post-hoc test (Tukey), */# p ≤ 0.05, **/## p ≤ 0.01, ***/### p ≤ 0.001, */**/*** relative to BSA+CF in (a-h) or BSA/Ctrl siR+CF (i-n), #/##/### relative to PA+CF (a-h) or PA/Ctrl siR+CF (i-n). Scale Bar: 25 µM in (e) and (g).

We next asked whether this pathway activation is functionally required for the PA+CF phenotype. Pharmacological inhibition of the core pyroptosis axis was achieved using MCC950 or VX-765, targeting NLRP3 or CASP1/4/5 (**Fig. 1d**). To functionally address potential changes in the inflammatory microenvironment, we analyzed the differentiation of fluorescently labelled monocytic THP1 cells into adherent macrophage-like cells. Thus, both inhibitors led to reduced PA-associated overactivation of monocytes during hPdLFs mechanoresponse, although without achieving complete normalization compared to the compressed BSA control (**Fig. 1e, f**). Since hyperinflammatory environments are also critical for the activation of osteoclasts (OCs) and, consequently, for bone resorption, we next analyzed OC activation by TRAP staining revealing partial normalization (**Fig. 1g, h**). To identify potentially relevant inflammasome/pyroptosis pathway components in PA+CF cultures, we performed an siRNA-mediated knockdown screen across eleven inflammasome– and pyroptosis-associated targets. This included sensors (NLRP3, AIM2, NLRC4), the adaptor ASC, caspases (CASP1, CASP4, CASP5) and gasdermins (GSDMA, GSDMB, GSDMD, and GSDME). When analyzing IL-6 levels (**Fig. 1i**), knockdown of *NLRP3*, *ASC*, and *CASP1* (canonical inflammasome axis), together with CASP4 (non-canonical) and GSDMD produced the strongest reductions in cytokine secretion. This was further accompanied by significant reductions in the PA-associated immune cell overactivation (**Fig. 1j, k**), at least with downregulation of *NLRP3*, *CASP1*, *CASP4* or *GSDMD*. Furthermore, the ratio of RANKL to OPG secretion (**Fig. 1l**) and osteoclast activation (**Fig. 1m, n**) were significantly reduced under these target-specific deficient conditions. Taken together, these results suggest that the inflammasome/pyroptosis signaling pathway contributes significantly, although only partially, to the PA-associated hyperinflammatory mechanoresponse of PdL cells.

### GDF15 is induced and relocalized under palmitic acid, contributing to the overactivated mechanoresponse

Since palmitate has recently been shown to modulate nuclear GDF15 levels in skeletal muscle [27], we next asked whether GDF15 might act as an upstream regulator of this pathway activation. GDF15 expression was increased in PA+CF compared to BSA+CF and OA+CF cultures (**Fig. 2a**). However, immunofluorescent staining revealed decreased nuclear GDF15 under PA+CF (**Fig. 2b, c**), with a corresponding increase in secreted GDF15 detected by ELISA in the culture supernatant (**Fig. 2d**). To determine whether GDF15 contributes to PA-associated hyperinflammatory mechanoresponse of PdL fibroblasts, siRNA-mediated downregulation was performed under hyperlipidemic conditions. Compared to control siRNA-treated PA+CF cultures, significantly reduced IL-6 secretion was observed in *GDF15*-deficient PA+CF cultures (**Fig. 2e**). This was further accompanied by a reduced overactivation of immune cells (**Fig. 2f, g**). However, the level of activated THP1 macrophages under GDF15 knockdown was still higher than observed in compressed BSA controls. Palmitic acid significantly shifted the RANKL/OPG ratio towards higher levels (**Fig. 2h**), accompanied by increased OC activation compared to compressed BSA controls (**Fig. 2i, j**). However, with *GDF15* deficiency, both the RANKL/OPG ratio and OC overactivation partially resembled the conditions observed in the BSA+CF group. Together, these data emphasize GDF15 as a partial contributor to the PA-associated hyperinflammatory and osteoclast-activating mechanoresponse of PdL fibroblasts.

**Figure 2:**
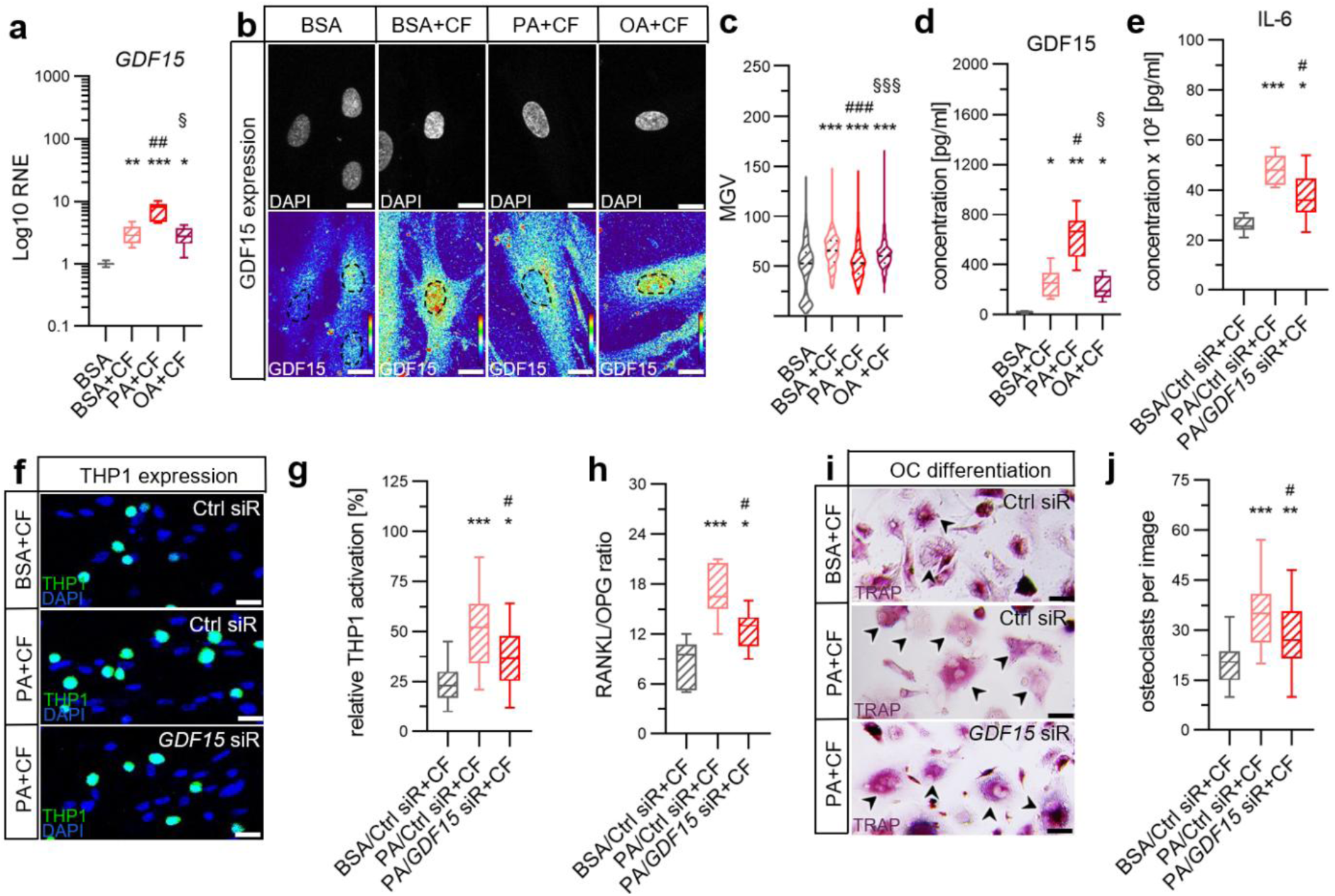
GDF15 is induced and relocalized toward the secreted route under palmitic acid, contributing to the overactivated mechanoresponse of PdL fibroblasts. **(a)** Relative normalize expression (RNE) of GDF15 in compressed PA and OA-treated hPdLFs compared to the compressed and unstressed BSA controls. Expression level is displayed in relation to the unstressed BSA control. **(b, c)** Immunofluorescent staining of nuclear GDF15 (thermal LUT, lower panel, b) with nuclei staining (DAPI, upper panel, b) and dotted line to visualize cell nuclei, quantified and displayed as mean grey value (MGV, c). **(d)** Secreted GDF15 levels in the supernatant. **(e)** IL-6 secretion following *GDF15* or control siRNA knockdown in PA-treated hPdLFs. **(f, g)** Monocyte activation following GDF15 knockdown. **(h)** RANKL/OPG ratio following GDF15 knockdown. (i, j) Osteoclast activation following GDF15 knockdown. One-way ANOVA with post-hoc test (Tukey), */#/§ p ≤ 0.05, **/## p ≤ 0.01, ***/###/§§§ p ≤ 0.001, */**/*** relative to BSA (a-e) or BSA/Ctrl siR+CF (f-j), #/##/### relative to BSA+CF (a-e) or PA/Ctrl siR+CF (f-j), §/§§§ relative to PA+CF (a-e). Scale Bar: 10 µM

### GDF15 knockdown partially reduces pyroptosis-associated caspase activity, alongside reduced apoptotic caspase activity

To determine whether the effect of GDF15 on the hyperinflammatory phenotype is mediated by the inflammasome/pyroptosis signaling pathway, we analyzed relevant pathway markers following GDF15 knockdown. Compared to control siRNA-treated PA+CF cultures, *GDF15*-deficient PA+CF cultures showed a modest reduction in IL-1β, IL-18, and HMGB1 secretion (**Fig. 3a**) and LDH release (**Fig. 3b**). This was further accompanied by a reduced activation of CASP1, but no changes in CASP4 activity (**Fig. 3c**). Additionally, *GDF15*-deficient PA+CF cultures showed reduced activity of the apoptosis-related CASP3 (**Fig. 3d**). To determine whether GDF15 acts additively with the canonical and non-canonical inflammasome axis, GDF15 was knocked down in combination with either CASP1 or CASP4 in PA+CF cultures. Compared to *GDF15* knockdown alone, both combinations further reduced immune cell (**Fig. 3e, f**) and osteoclast activation (**Fig. 3g, h**). Together, these data indicate that GDF15’s contribution to pyroptosis-associated hyperinflammation is partial and additive to the inflammasome axis.

**Figure 3:**
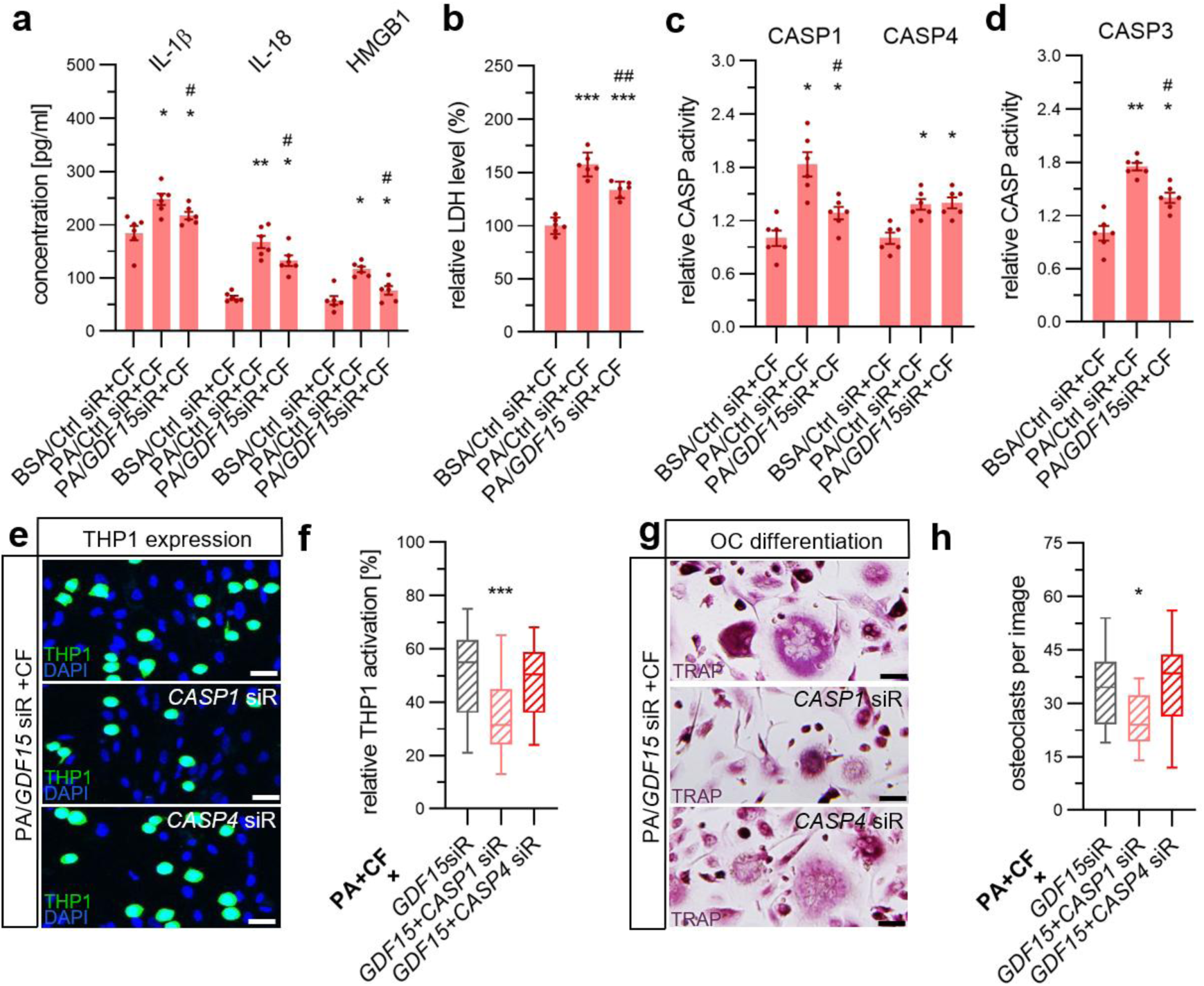
GDF15 knockdown partially reduces pyroptosis-associated hyperinflammation. **(a)** Secreted IL-1β, IL-18, and HMGB1 in compressed PA-treated hPdLFs after *GDF15* or control siRNA knockdown **(b)** LDH release **(c)** CASP1 and CASP4 activity. **(d)** CASP3 activity. **(e, f)** Monocyte activation following combined GDF15/CASP1 or GDF15/CASP4 knockdown in compressed PA-treated hPdLFs. **(g, h)** osteoclast activation. One-way ANOVA with post-hoc test (Tukey), */# p ≤ 0.05, **/## p ≤ 0.01, *** p ≤ 0.001, */** relative to BSA/Ctrl siR+CF (a-d) or PA/GDF15 siR+CF (e-h), #/## relative to PA/Ctrl siR+CF.

### GDF15 signals through extracellular and intracrine, nuclear-export-dependent routes modulating excessive mechanoresponses

Next, we aimed to investigate by which signaling pathway GDF15 modulates the mechanoresponse of PdL cells. ALK1, ALK2, and ALK5 have previously been identified as GDF15-binding receptors expressed in hPdLFs [18]. Following this, we investigated the functional relevance of the extracellular, receptor-mediated signaling pathway using the receptor inhibitors MU1700 (ALK1/ALK2) and TP-008 (ALK5) (**Fig. 4a**). Compared to untreated PA+CF cultures, receptor blockade was observed to partially decrease the overactivation of immune cells (**Fig. 4b, c**) and osteoclasts (**Fig. 4d, e**), although the phenotype was not completely eliminated. This may indicate an additional, receptor-independent signaling pathway for GDF15.

**Figure 4:**
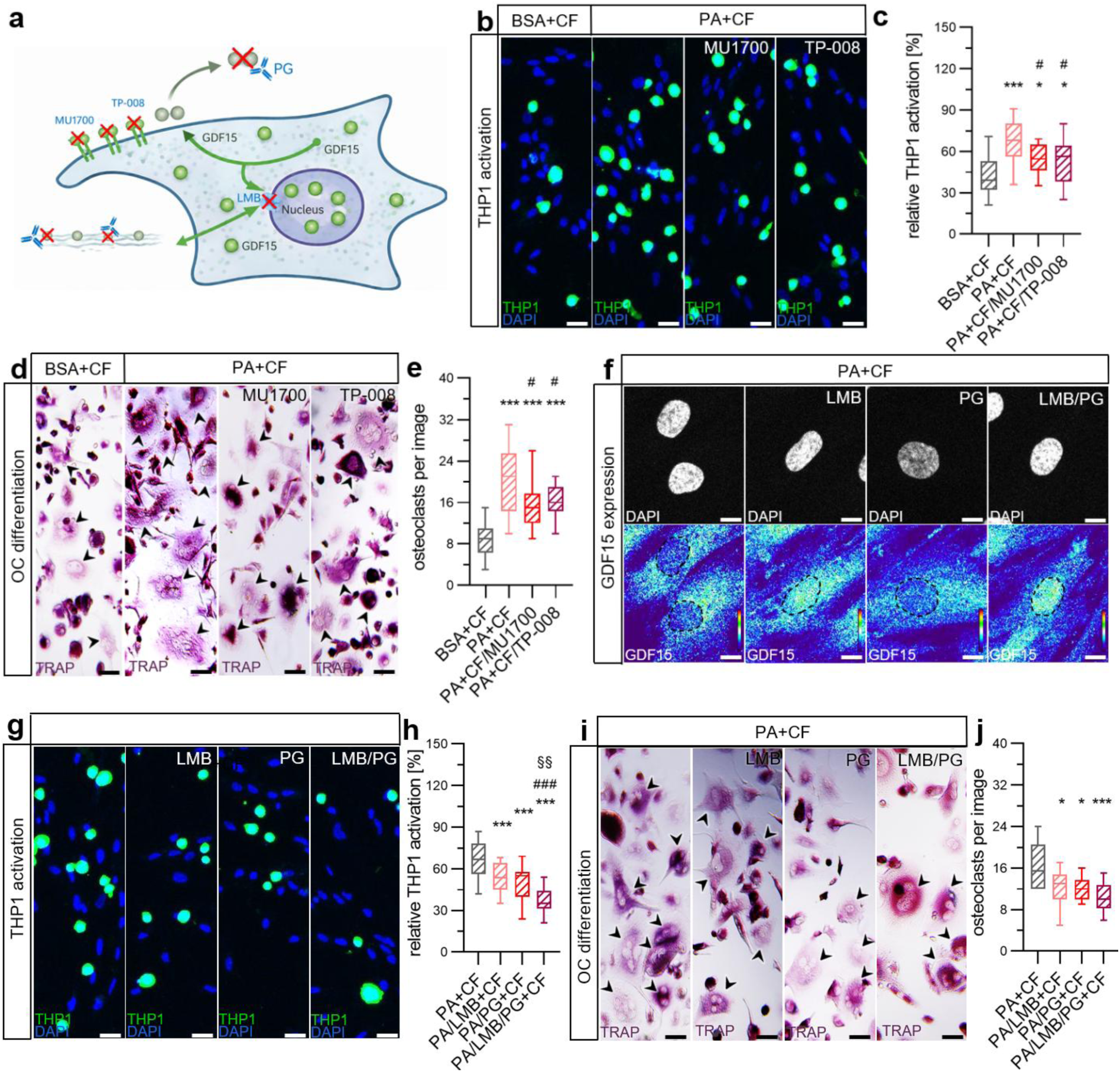
Specific GDF15 signaling routes partially mediating the PA+CF hyperinflammatory mechanoresponse. **(a)** Schematic of ALK1/ALK2 blockade (MU1700), ALK5 blockade (TP-008), nuclear export inhibition by leptomycin B (LMB), and extracellular GDF15 blockade (Ponsegromab, PG). **(b, c)** Monocyte activation under MU1700 or TP-008 treatment of compressed PA-treated hPdLFs. (**d, e**) osteoclast activation under MU1700 or TP-008 treatment of compressed PA-treated hPdLFs. **(f)** Immunofluorescent staining of nuclear GDF15 (thermal LUT, lower panel) following LMB, PG, or combined LMB+PG treatment of compressed PA treated hPdLFs with nuclei staining (DAPI, upper panel) and dotted line to visualize cell nuclei. (**g, h**) Monocyte activation. (**i, j**) osteoclast activation. One-way ANOVA with post-hoc test (Tukey), */# p ≤ 0.05, §§ p ≤ 0.01, ***/### p ≤ 0.001, */*** relative to BSA+CF (b-e) or PA+CF (g-j), #/### relative to PA+CF (b-e) or PA/LMB+CF (h), § in relation to PA/PG+CF (h).

To address this, we investigated the intracrine, nuclear route of GDF15 signaling using Leptomycin B (LMB) to block nuclear export and Ponsegromab (PG), a monoclonal GDF15-targeting antibody, to block extracellular GDF15 (Fig. 4a). Immunofluorescent staining showed increased nuclear GDF15 in PA+CF cultures treated with LMB, alone or in combination with PG, compared to untreated PA+CF cultures (Fig. 4f). We next asked whether this shift in GDF15 localization induced by LMB translates into a functional change of immune cell and osteoclast overactivation. Compared to untreated PA+CF cultures, LMB treatment reduced the overactivation of both immune cells (**Fig. 4g, h**) and osteoclasts (**Fig. 4i, j**), and blocking extracellular GDF15 with Ponsegromab resulted in a comparable reduction in both parameters. Combining LMB and PG promoted the strongest effect on monocyte activation, exceeding either treatment alone. Together, these data indicate that GDF15 acts through both extracellular and intracrine signaling routes to partially mediate the PA-associated hyperinflammatory phenotype.

## Discussion

Orthodontic tooth movement is based on a controlled, force-induced inflammatory response in the periodontal ligament, which is subsequently accompanied by tissue and alveolar bone remodeling [1]. Excessive mechanoresponses of the PdL increase the probability of treatment-associated risks including root resorption and tooth loss [9]. Obesity-associated hyperlipidemia was reported to induce hyperinflammatory PdL responses under mechanical stress with the potential triggers and mechanisms appearing to be highly tissue-specific and multifactorial [6]. This study now demonstrated that hyperlipidemia, specifically by saturated fatty acids such as palmitic acid, overactivates the inflammasome/pyroptosis pathway in mechanically stressed PdL cells, contributing to their hyperinflammatory and osteoclast-activating mechanoresponse. The TGF-β/BMP superfamily member GDF15 was identified as a partial regulator of this response, signaling through both extracellular and intracrine routes.

Compressive forces during orthodontic therapy were demonstrated to activate the inflammasomal pathway potentially leading to pyroptosis in the PdL [28, 29]. In this study, hyperlipidemic conditioning with palmitic acid, but not oleic acid, further increased the pathway activation *in vitro* with enhanced activation of relevant caspases including CASP1 and CASP4, increased secretion levels of specific cytokines including IL-1β and IL-18, as well as HMGB1 and extracellular release of LDH. This matches the established concept that saturated, but not unsaturated, fatty acids can prime and activate the NLRP3 inflammasome [11, 12, 30]. It is also consistent with a recent study of Sun et al. [13] demonstrating that palmitate specifically induces PdL cell pyroptosis via a caspase-4/GSDMD-mediated pathway. Pharmacological inhibition of the pyroptosis pathway with both the NLRP3-specific inhibitor MCC950, and CASP1/4/5-inhibiting VX-765 balanced the mechanoresponse under palmitic acid overexposure confirming that the pathway is functionally required for the full mechanoresponse. A plausible mechanism for this might involve the pyroptosis-associated release of IL-1β, as it could then act on remaining PdL cells through its receptor IL1R1 to promote a pro-inflammatory/resorptive cellular phenotype, including increased production of IL-6 and RANKL [31–33]. A supporting population-level loop has been already described for the PdL in connection to periodontitis [34]. Mo et al. [34] demonstrated that distinct fibroblast subpopulations in the PdL produce IL-1β and RANKL respectively, with IL-1β acting in a paracrine manner on the RANKL-producing population. There, IL-1β promoted an osteoclast-activating output, alongside elevated cleaved caspase-1 and NLRP3 in periodontitis tissue *in vivo*. Pyroptosis of a subset of PdL cells may therefore contribute to activating a broader response in the surviving population, though this remains speculative and untested in relation to the present findings. However, siRNA screen across eleven inflammasome– and pyroptosis-associated targets pointed to CASP1, in addition to CASP4, as particularly important regulators of this functional mechanoresponse-associated output. Knockdown of *CASP1* and *CASP4* reduced both immune cell and osteoclast overactivation, alongside *NLRP3* and *GSDMD* downregulation. This is partially in contrast to the results of Sun et al. [13], where CASP4 and GSDMD, rather than CASP1, were identified as the drivers of palmitate-induced pyroptotic cell death. However, this study was performed without additional compression and with a substantially higher palmitate concentration of 500 µM over a much shorter exposure of 24 hours, closer to an acute, high-dose experimental model. Even though there appear to be mechanistic differences between these models, the present findings overall confirm the relevance of the inflammasome/pyroptosis pathway for the regulation of inflammation and bone remodeling in the PdL.

GDF15 was investigated as a potential upstream modulator of this pathway, given its established role in the regulation of immune cell and osteoclast activation in the PdL mechanoresponse [16–21, 35]. In this study, GDF15 expression was increased in compressed PdL fibroblasts under palmitic acid exposure. Nuclear GDF15 levels were decreased, while GDF15 secretion was increased, a pattern potentially consistent with a shift in subcellular distribution toward the secreted route. This matches reports that palmitate reduces nuclear GDF15 in skeletal muscle through a SMAD3-PAI1-dependent mechanism [27]. Two partly independent stress pathways make this shift plausible in the present model. One is an endoplasmic reticulum stress route acting through PERK, eIF2, and CHOP, which directly binds the *GDF15* promoter, shown in PMA-differentiated THP1 cells and primary monocyte-derived macrophages [36]. The study demonstrated a specific impact of saturated but not unsaturated fatty acids. The other is a lysosomal route in which palmitic acid triggers lysosomal calcium release and TFEB nuclear translocation, driving GDF15 transcription as part of a protective lysosomal stress response in adipose tissue macrophages [37]. In an animal study, where mice received fatty-acid-enriched diets, palmitic acid enrichment positively correlated with serum GDF15 levels 8 hours after feeding [38]. Interestingly, also for other fatty acid enrichments, including oleic acid, a comparable correlation was also observed even at an earlier analytical time point. This delay may reflect differences in intestinal absorption or metabolic processing between fatty acid types, though the precise mechanism was not addressed in that study. However, a direct comparison of these short-term systemic kinetics with the present *in vitro* findings, based on local, cell-autonomous production over a considerably longer exposure period, is anyhow only possible to a limited extent. Several factors may account for this, which include differences in the fatty acid concentrations used. Elevated serum palmitic acid is a recognized feature of human obesity and hyperlipidemia [39], and the concentration applied here was chosen accordingly, though it may not directly correspond to the circulating fatty acid levels achieved in the mouse study. Interspecies differences in fatty acid metabolism may further limit the comparability of the two models. Moreover, the cellular context also differs, since PdL fibroblasts may not be the main GDF15-secreting cell type under hyperlipidemic conditions. Besides this, GDF15 protein dynamics add a further layer of complexity, including its short serum half-life of about 3 hours [40], and its various, potentially cell– and context-dependent modes of action through local, autocrine, and paracrine signaling [41–43].

*GDF15* knockdown reduced immune cell and osteoclast overactivation during mechanoresponse of PdL fibroblasts under palmitic acid exposure. This is consistent with its established role in the PdL mechanoresponse under other stress conditions, where its knockdown or blockade similarly reduced immune cell activation and RANKL/OPG-associated osteoclast activity [17–21, 35]. Outside the periodontal ligament, the role of GDF15 in immune cell regulation is rather described as anti-inflammatory and immunosuppressive, dampening macrophage, neutrophil, dendritic cell, natural killer cell, and T lymphocyte activity [44, 45]. Its impact on osteoclast regulation is less consistent, with reports of both pro– and anti-osteoclastogenic effects depending on cellular context and stimulus [46, 47].

GDF15 deficiency reduced the inflammasome/pyroptosis pathway activation in compressed hyperlipidemic PdL fibroblasts, reflected in decreased secretion of IL-1β, IL-18, HMGB1, and LDH. However, palmitic acid-associated changes in GDF15 signaling only partially affected this pathway and downstream functional outputs as demonstrated by the combined knockdown with CASP1 and CASP4. The inflammasome/pyroptosis pathway has been previously linked to GDF15 in the opposite direction, with GDF15 shown to suppress NF-κB and inflammasome signaling in macrophages and other myeloid cells [48, 49]. The present findings in PdL fibroblasts may therefore provide first evidence of a promoting role in a mesenchymal cell context. Since GDF15 signals through cell type-specific receptor complexes [18, 50, 51], a mesenchymal-versus-myeloid difference in downstream signaling offers a plausible explanation.

Compressed PdL fibroblasts also showed increased CASP3 activation levels when additionally exposed to palmitic acid. This CASP3 overactivation was reduced by GDF15 knockdown, indicating a role for this cytokine in stress-induced apoptosis regulation alongside the impact on pyroptosis. It is well established that mechanical stress leads to upregulation of cell death programs in PdL cells [52–54]. A comparable promoting role was demonstrated for palmitic acid in various cell types [55, 56]. The observed impact of GDF15 further matches reports of reduced ER-stress-induced apoptosis after GDF15 knockdown in pancreatic beta cells [57]. However, exogenous stimulation with GDF15 instead reduced caspase-3/7 activity in oral squamous cell carcinoma [58], which further highlights the relevance and difference of specific GDF15 signaling routes.

GDF15 exists in two forms with distinct locations and functions. In its uncleaved precursor form, GDF15 is located intracellularly and can be transported into the nucleus, where it modulates gene transcription [22]. This precursor form can also be secreted and bound to structural components of the ECM, forming local stromal stores. The cleaved, mature form, by contrast, is released freely into circulation [23]. PdL fibroblasts express GDF15-binding receptors of the ALK family [18]. Blocking ALK1, ALK2, or ALK5 reduced the overactivation of immune cells and osteoclasts in compressed, hyperlipidemic PdL fibroblasts in this study. In human airway epithelial cells, GDF15 has been reported to bind ALK1, driving cellular senescence [59]. In dorsal root ganglion sensory neurons, GDF15 signals through ALK2, inhibiting Nav1.8 sodium channel activity [60]. ALK5 forms a receptor heterodimer with TGF-βRII through which GDF15 signals in myeloid cells, demonstrated directly on THP-1 cells [50]. An autocrine activation loop may exist, in which fibroblast-secreted GDF15 binds ALK receptors on the fibroblasts themselves, contributing to the mechanoresponse under hyperlipidemic conditions. The comparable effect of Ponsegromab, a GDF15-specific monoclonal antibody currently in phase II trials, is consistent with this. Since THP-1 cells were also shown to express ALK5 [50], GDF15 may additionally act on them directly, altering their macrophage and osteoclastogenic differentiation in this study. This is in line with Li et al., who demonstrated that recombinant GDF15 protein directly stimulates the differentiation of osteoclast precursors, including THP-1-derived macrophages and RAW264.7 cells [35]. Beyond receptor-mediated signaling at the cell surface, GDF15 may also act through an intracrine, nuclear route. Pro-GDF15 accumulates in the nucleus and interrupts Smad-dependent transcription, demonstrated across several cell lines including osteosarcoma, colorectal cancer, and lung adenocarcinoma cells [22]. Whether this nuclear route shapes the PdL fibroblast’s own transcriptional program, and thereby its secretion of the pro-inflammatory and osteoclastogenic factors measured in this study, remains untested and offers a candidate mechanism for future investigation.

Taken together, pyroptosis and GDF15 are each partial, non-exclusive contributors to a hyperinflammatory mechanoresponse of PdL fibroblasts under hyperlipidemia. However, several limitations of this study should be acknowledged. This in vitro model does not fully replicate the in vivo complexity of the periodontal ligament. Palmitic acid concentration, force magnitude, and exposure duration were chosen to reflect clinically relevant conditions, but only a single combination was tested. Donor age, sex, and health status may further influence PdL fibroblast behavior [61, 62]. Despite these limitations, the findings carry clinical relevance. Through the RANKL/OPG-osteoclast axis discussed above, this response carries a potential risk for root resorption and tooth loss in patients with hyperlipidemia undergoing orthodontic treatment. Blocking pyroptosis in a closely related model reduced root resorption without impairing tooth movement itself [28], indicating that the pathological, excessive component of this mechanoresponse might be targeted.

## Acknowledgment

We thank Dr. Daniela Korth and Sushil Kumar Dubey for excellent technical assistance. We thank Dr. Sebastian Drube for valuable advice on inflammation-related aspects of the study and for technical support. We further thank Prof. Dr. Christian Hübner for supporting confocal microscopic imaging using the LSM 880.

## Author contributions

M.B.: Investigation, Writing – original draft, Visualization. A.M.: Methodology, Validation, Investigation. F.G.Z.: Investigation. R.Y.: Investigation. A.D.: Investigation. C.-L.H.: Investigation. U.S.-S.: Conceptualization, Methodology, Writing – review & editing. J.S.: Conceptualization, Methodology, Investigation, Writing – review & editing, Visualization, Supervision, Project administration. C.J.: Conceptualization, Writing – review & editing, Supervision, Project administration, Funding acquisition.

## Generative AI statement

During the preparation of this manuscript, the authors used Claude (Anthropic, San Francisco, CA, USA) to assist with language refinement. All AI-assisted content was critically reviewed and edited by the authors, who take full responsibility for the final content of the manuscript.

## Data availability

The data generated and analyzed during the current study are available from the corresponding author upon reasonable request.

